# Misoprostol Attenuates Cardiomyocyte Proliferation in the Neonatal Heart through Bnip3 and Perinuclear Calcium Signaling

**DOI:** 10.1101/681692

**Authors:** Matthew D. Martens, Jared T. Field, Nivedita Seshadri, Chelsea Day, Donald Chapman, Richard Keijzer, Christine R. Doucette, Grant M. Hatch, Adrian R. West, Tammy L. Ivanco, Joseph W. Gordon

## Abstract

Systemic hypoxia resulting from preterm birth, altered lung development, and cyanotic congenital heart disease is known to impede the regulatory and developmental pathways in the neonatal heart. While the molecular mechanisms are still unknown, hypoxia induces aberrant cardiomyocyte proliferation, which may be initially adaptive, but can ultimately program the heart to fail in early life. Recent evidence suggests that the prostaglandin E1 analogue, misoprostol, is cytoprotective in the hypoxia-exposed neonatal heart by impacting alternative splicing of the Bcl-2 family member Bnip3, resulting in the generation of a variant lacking the third exon (Bnip3**Δ**Exon3 or small Nip; sNip). Using a rodent model of neonatal hypoxia, in combination with rat primary neonatal cardiomyocytes (PVNCs) and H9c2 cells, we sought to determine if misoprostol can prevent cardiomyocyte proliferation and what the key molecular mechanisms might be in this pathway. In PVNCs, exposure to 10% oxygen induced myocyte proliferation concurrent with molecular markers of cell-cycle progression, such as Cyclin-D1, which were prevented by misoprostol treatment. Furthermore, we describe a critical role for sNip in opposing cardiomyocyte proliferation through several mechanisms, including reduced expression of the proliferative MEF2C-myocardin-BMP10 pathway, accumulation of nuclear calcium leading to NFATc3 activation, and increased expression of the cardiac maturation factor BMP2. Intriguingly, misoprostol and sNip inhibited hypoxia-induced glycolytic flux, which directly influenced myocyte proliferation. These observations were further supported by knockdown studies, where hypoxia-induced cardiomyocyte proliferation is restored in misoprostol-treated cells by an siRNA targeting sNip. Finally, in postnatal day (PND)-10 rat pups exposed to hypoxia, we observed histological evidence of increased nuclei number and increased PPH3 staining, which were completely attenuated by misoprostol treatment. Collectively, this data demonstrates how neonatal cardiomyocyte proliferation can be pharmacologically modulated by misoprostol treatment, which may have important implications for both neonatal and regenerative medicine.

## Introduction

Systemic neonatal hypoxia resulting from early-life cyanotic events including preterm birth, placental abnormalities, abnormal/impaired lung development, and certain forms of congenital heart disease, are known drivers of both structural and functional changes in the developing heart [1,2]. Specifically, these infants demonstrate dramatic reductions in systolic and diastolic function, concurrent with extensive increases in left-ventricular mass [3]. While such changes are thought to be initially protective to preserve cardiac output in the neonatal stage, they also significantly increase the risk for early-life heart failure, and lifelong cardiovascular abnormalities [1,3,4].

Although traditional literature suggests that cardiomyocytes proliferate early in gestation, and that postnatal cardiac growth is achieved exclusively through hypertrophy, recent evidence in humans demonstrates that cardiomyocytes continue to proliferate in the first year of life, and a small number remain proliferative for as long as 20 years [5]. Moreover in rodents, cardiomyocytes retain the ability to proliferate until the point of binucleation, which occurs on or before postnatal day 7 [6,7]. Work focused specifically on this developmental window has revealed that cardiomyocytes continue to undergo mitosis in an environment where the oxygen tension is low, even extending into adulthood, where whole animal hypoxia-exposure following myocardial infarction reduces cardiac damage and fibrosis by enhancing cardiomyocyte proliferation [5,8,9]. In addition, sustained cardiomyocyte proliferation elicited by hypoxia is also dependent on a metabolic shift towards enhanced glycolysis, and impaired mitochondrial respiration [10,11]. Importantly, infants exposed to neonatal hypoxia display evidence of smaller and more numerous myocytes, suggesting that cardiac remodelling occuring at this age is substantially different from the remodelling occurring in adulthood [11].

The developmental switch from proliferation to hypertrophy is the result of large-scale shifts in gene expression and metabolism, where changes in the expression pattern of the myocyte enhancer factor-2 (MEF2) family appears to be a key event [12–15]. Genetic studies have shown that three MEF2 genes have non-redundant and potentially antagonistic roles. For example, MEF2C has been shown to regulate cardiomyocyte proliferation during cardiac looping, MEF2A promotes cardiomyocyte differentiation and mitochondrial biogenesis, and MEF2D modulates postnatal maturation and remodelling [13–17]. Furthermore, MEF2C expression is governed by hypoxia signalling, where loss of the hypoxia-inducible factor-1α (HIF1α) prevents MEF2C expression, cardiac looping and embryo viability [18,19]. Myocardin is a transcriptional co-activator involved in the proliferation, differentiation and maturation of cardiomyocytes, and is a direct transcriptional target of MEF2C [13,16,17,19]. Myocardin’s ability to regulate cardiomyocyte proliferation is directly linked to the expression of bone morphogenetic protein (BMP) 10, a growth factor associated with embryonic cardiomyocyte expansion [2,20–22].

In contrast to proliferation, cardiomyocyte hypertrophy signaling is largely dominated by the calcium-calmodulin-dependent phosphatase, calcineurin (PP2B) [23–26]. Classically, calcineurin is activated by low amplitude nuclear calcium transients, resulting in nuclear accumulation of nuclear factor of activated T-Cells (NFAT) [23,27]. In the nucleus, NFATs complex with GATA4 to drive the expression of many hypertrophy genes including beta-myosin heavy chain (β-MHC), brain natriuretic peptide (BNP), myocyte-enriched calcineurin-interacting protein 1 (MCIP1), and endothelin 1 (ET-1) [Reviewed by Wilkins and Molkentin, 2002 [25]]. In addition to its interactions with GATA4, NFAT also physically interacts with the p65 subunit of nuclear factor kappa B (NF-κB) and is able to drive differentiation and hypertrophy (12), and increase the expression of the cardiac maturation factor BMP2 [28]. Calvienurin has additionally been shown to enhance MEF2A activity [29,30].

Independent from the hypoxia-inducibility of MEF2C, HIF1α also drives the expression of BCL2/Adenovirus E1B 19 KDa Protein-Interacting Protein 3 (Bnip3), whose protein products play a pivotal role in hypoxia-induced cardiomyocyte apoptosis, necrosis and autophagy [31–37]. Alternative splicing of Bnip3 produces a truncated form of the protein (ie. short/small Nip or sNip) lacking the second exon in humans, or the third exon in rodents, that acts as an endogenous inhibitor of the full-length protein [20,38]. Recently, we have shown that these spliced variants also drive the accumulation of calcium in the nucleus triggering hypertrophic growth of cardiomyocytes. Interestingly, we also demonstrated that the prostaglandin analogue, misoprostol can modulate the expression of sNip in multiple cell and tissue types [20,39].

Building on this previous work, in this report we provide evidence that misoprostol inhibits hypoxia-induced proliferation in the neonatal heart, primary ventricular neonatal cardiomyocytes and cultured cell lines by reducing MEF2C and BMP10 expression. We further show that hypoxia in combination with misoprostol treatment enhances sNip expression leading to nuclear calcium retention, activation of NFATc3, and increased BMP2 expression to promote maturation and hypertrophy. Finally, sNip-dependent NFATc3 activation prevents hypoxia-induced glycolytic flux, which directly opposes proliferation.

## Materials and Methods

### In vivo neonatal hypoxia model

All procedures in this study were approved by the Animal Care Committee of the University of Manitoba, which adheres to the principles for biomedical research involving animals developed by the Canadian Council on Animal Care (CCAC). Litters of Long-Evans rat pups and their dams were placed in a hypoxia chamber with 10% O_2_ (±1%) from postnatal day (PND) 5-10. Additionally, hypoxic animals were exposed to 100% N_2_ for 90 seconds, daily from PND 5-7 (inclusive). Control litters (N=3), were left in normoxic conditions at 21% O_2_. Animals received 10 μg/kg misoprostol or saline control, administered orally in rat milk substitute daily from PND5-10. At PND10 animals were euthanized and perfused with saline for tissue collection.

### Histology

Male PND10 hearts were fixed in 10% buffered formalin for 24 hours at time of sacrifice and stored in phosphate buffered saline (PBS; Hyclone). Fixed hearts were cut longitudinally (to expose the four chambers of the heart), processed and embedded in paraffin blocks. Hearts were sectioned at 5μm thicknesses, mounted on APTES coated slides, and stained with Hematoxylin and Eosin, Masson’s Trichrome, or TUNEL via the CardioTACS™ In Situ Apoptosis Detection Kit (Trevigen) as per the manufacturer’s protocol by the University of Manitoba Histology Core. Hearts were imaged on a Zeiss Axio Lab.A1 bright-field microscope fitted with an Axiocam 105 color camera (Zeiss). Imaging was done using Zen 2.3 Pro imaging software and quantification, scale bars, and processing was done on Fiji (ImageJ) software.

### Plasmids and virus production

The endoplasmic reticulum (CMV-ER-LAR-GECO1), and nuclear (CMV-NLS-R-GECO) targeted calcium biosensors were gifts from Robert Campbell (Addgene #61244, and 32462) [40,41]. CMV-dsRed was a gift from John C. McDermott and myc-NFATc3 was a gift from Tetsuaki Miyake. GW1-Peredox-mCherry (Peredox-mCherry) was a gift from Gary Yellen (Addgene plasmid #32380) [42]. The FRET-based lactate biosensor [Laconic/pcDNA3.1(-)] and pyruvate biosensor [Pyronic/pcDNA3.1(-)] were gifts from Luis Felipe Barros (Addgene plasmid #44238, and #51308) [43,44]. The mouse HA-Bnip3ΔExon3 (sNip) (Accession #MF156210) were described previously (Addgene #100793) [39]. Generation of the pLenti-Bnip3ΔExon3 (sNip) virus using a pLenti-puro back bone, which was described previously [20].

### Cell culture, transduction and transfections

Rat primary ventricular neonatal cardiomyocytes (PVNC) were isolated from 1-2-day old pups using the Pierce Primary Cardiomyocyte Isolation Kit (#88281). H9c2 cells were maintained in Dulbecco’s modified Eagle’s medium (DMEM; Hyclone), containing penicillin, streptomycin, and 10% fetal bovine serum (Hyclone), at 37 °C and 5% CO2. All cells were transfected using JetPrime Polyplus reagent, as per the manufacturer’s protocol. Lentiviral expression of sNip and RNAi targeting of sNip were described previously [20]. For misoprostol treatments, 10 mM misoprostol (Sigma) in phosphate buffered saline (PBS; Hyclone) was diluted to 10 μM directly in media and applied to cells for 48 hours. For hypoxia treatments, cells were held in a Biospherix incubator sub-chamber with 10% O_2_ (±1%), 5% CO_2_, balanced with pure N_2_ (regulated by a Biospherix ProOx C21 sub-chamber controller) at 37 °C for 48 hours. 2-aminoethoxydipheny borate (2APB), etomoxir (ETO), Cyclosporine-A (CsA) and 2-Deoxy-D-Glucose (2DG) were purchased from Sigma.

### Fluorescent staining, live cell imaging and immunofluorescence

Hoechst 3334 and Calcein-AM were all purchased from Biotium and applied using the manufacturer’s protocol. Tag-it Violet Proliferation and Cell Tracking Dye was purchased from BioLegend and applied using the manufacturer’s protocol. Glucose uptake assay was measured as previously described using the fluorescent D-glucose analog 2NBDG (200 μM; Molecular Probes) [15]. Immunofluorescence with Myc-Tag (CST # 2272) and Phospho-Histone H3 (CST # 9701) antibodies were performed in conjunction with fluorescent secondary antibodies conjugated to Alexa Fluor 466 or 594 (Jackson) in fixed and permeabilized PND10 heart tissues, PVNCs and H9c2’s. All imaging experiments were done on a Zeiss Axiovert 200 inverted microscope fitted with a Calibri 7 LED Light Source (Zeiss) and Axiocam 702 mono camera (Zeiss), while FRET imaging was done using a Cytation 5 Cell Imaging Multi-Mode Reader. Imaging was done using Zen 2.3 Pro or Gen5 imaging software and quantification, scale bars, and processing including background subtraction, was done on Fiji (ImageJ) software.

### Measurement of Proliferation by Flow Cytometry

Cell cycle was assessed by flow cytometry using the Nicoletti method and FxCycle™PI/RNase Staining Solution purchased from Invitrogen (product # F10797) [45]. Red fluorescence was analyzed for 20,000 cells/treatment using a Thermo Scientific Attune NxT flow cytometer equipped with a 488 nm laser, as per the manufacturer’s protocol. Gating and DNA histogram analysis was described previously [46].

### Immunoblotting

Protein isolation and quantification was performed as described previously [20]. Extracts were resolved via SDS-PAGE and later transferred to a PVDF membrane using an overnight transfer system. Immunoblotting was carried out using primary antibodies in 5% powdered milk or BSA (as per manufacturer’s instructions) dissolved in TBST. Horseradish peroxidase-conjugated secondary antibodies (Jackson ImmunoResearch Laboratories; 1:5000) were used in combination with enhanced chemiluminescence (ECL) to visualize bands. The following antibodies were used: Cyclin-D1 (sc-450), p57(KP39) (sc-56341), LDHA (CST #3582), MEF2C (CST #5030), MEF2A (CST #9736), Myc-Tag (CST # 2272), NFATc3 (F-1) (sc-8405), p-NFATc3 (Ser 169) (sc-68701), Histone-H3 (CST #4499), MEK1/2 (CST #8727), Actin (sc-1616), and Tubulin (CST #86298). For detection of the Bnip3 splice variant, Bnip3ΔExon3 (sNip), we used a custom rabbit polyclonal antibody that was previously described and validated [20].

### Extracellular Acidification

Extracellular acidification was determined on a Seahorse XF-24 Extracellular Flux Analyzer in combination with Seahorse Glycolysis Stress Test. Calculated acidification rates were determined as per manufacturer’s instructions (Glycolysis Stress Test; Seahorse).

### Real-Time PCR

Total RNA was extracted from cells and pulverized tissues by the TRIzol extraction method and genomic DNA was removed via the RNeasy Mini Kit (Qiagen), including an On-Colum DNase Digestion (Qiagen). For qRT-PCR, mRNA was extracted from cells using Trizol then reverse transcribed into cDNA. Following DNase treatment, cDNA was combined with SYBR Green Supermix (Thermo) and mRNA was amplified using the ABI 7500 Real-Time PCR system (Applied Biosystems). Primers were: MEF2C Mus Fwd: GATGCAGACGATTCAGTAGGTC, MEF2C Mus Rev: GGATGGTAACTGGCATCTCAA, MEF2A Rat FWD: GGAACCGACAGGTGACTTTTA, MEF2A Rat REV: AGAGCTGTTGAAGATGATGAGTG, BMP10 Rat Fwd: TGCCATCTGCTAACATCATCC, BMP10 Rat Rev: CAAACGATCTCTCTGCACCA, BMP2 Rat FWD: GCTCAGCTTCCATCACGAA, BMP2 Rat REV: GAAGAAGCGTCGGGAAGTT, MYOCD Rat Fwd: GTGTGGAGTCCTCAGGTCAAAC, MYOCD Rat Rev: TGATGTGTTGCGGGCTCTT, L13 Rat Fwd: AGGAGGCGAAACAAATCCAC, L13 Rat Rev: TATGAGCTTGGAGCGGTACTC.

### Statistics

Data are presented as mean ± standard error (S.E.M.). Differences between groups in imaging experiments with only 2 conditions were analyzed using an unpaired t-test, where (*) indicates P<0.05 compared with control. Experiments with 4 or more conditions were analyzed using a 1-way ANOVA, or 2-way ANOVA where appropriate, with subsequent Tukey test for multiple comparisons, where (*) indicates P<0.05 compared with control, and (**) indicates P<0.05 compared with treatment. All statistical analysis was done using GraphPad Prism software.

## Results

### Misoprostol opposes hypoxia-induced neonatal cardiomyocyte proliferation

In order to investigate whether misoprostol can modulate hypoxia-induced cardiomyocyte proliferation, we employed cultured primary ventricular neonatal cardiomyocytes (PVNCs), isolated from PND 1-2 rats pups. When exposed to 10% O_2_ for 48-hours, we observed a marked increase in cyclin-D1 expression, a protein that rapidly accumulates in the G_**1**_ phase of the cell cycle and is further required for the transition from the G_1_ to S phase (Fig. 1A). Furthermore, the induction of cyclin-D1 was completely absent with the addition of misoprostol. Using live cell imaging and a tracker-based proliferation technique, we observed that hypoxia significantly increased the number of proliferative PVNCs compared to control, which was inhibited with misoprostol treatment (Fig. 1B, -C). To confirm that hypoxia-induced proliferation was also happening at the cell population level, we utilized flow cytometry-based nicoletti assays with propidium iodide, and observed a significant increase in the ratio of G_2_ phase cells to G_1_ phase cells when cultures were exposed to 10% O_2_ for 48 hours, indicating a hypoxia-induced increase in mitosis (Fig. 1D, -E). However, when cells were treated with misoprostol and exposed to hypoxia, the ratio of G_2_ to G_1_ phase cells was reduced to control levels (Fig. 1D, -E). Next, we evaluated cell size by calculating surface area of PVNCs exposed to hypoxia and/or misoprostol (Fig. 1F, -G). Consistent with our previous results [20], misoprostol treatment increased the size of PVNCs, while exposure to hypoxia decreased cell area. However, when hypoxic PVNCs were treated with misoprostol, cell area was significantly restored. As fetal cardiomyocyte proliferation is associated with high glycolytic rates, and cardiomyocyte maturation involves considerable mitochondrial biogenesis and increased fatty acid oxidation, we evaluated whether inhibition of glycolysis with 2-deoxyglucose (2DG) or inhibition mitochondrial fatty acid uptake with etomoxir (ETO) altered the myocyte response to either hypoxia or misoprostol treatment. Shown in Figure 1H and -I, hypoxia-induced proliferation was attenuated in the presence of 2DG, and had no impact on the ability of misoprostol to block PVNC proliferation. Finally, etomoxir treatment of PVNCs increased proliferation in normoxic conditions, but did not enhance the effect of hypoxia (Fig. 1J). Intriguingly, misoprostol treatment was able to block hypoxia-induced proliferation, but not in the presence of etomoxir. Collectively, these observations suggest that hypoxia-induced proliferation is dependent on a switch to glycolysis, and that mitochondrial fatty acid oxidation may serve to oppose neonatal myocyte proliferation. These results further indicate that misoprostol’s ability to modulate proliferation is dependent on myocyte energetics.

**Figure 1.**
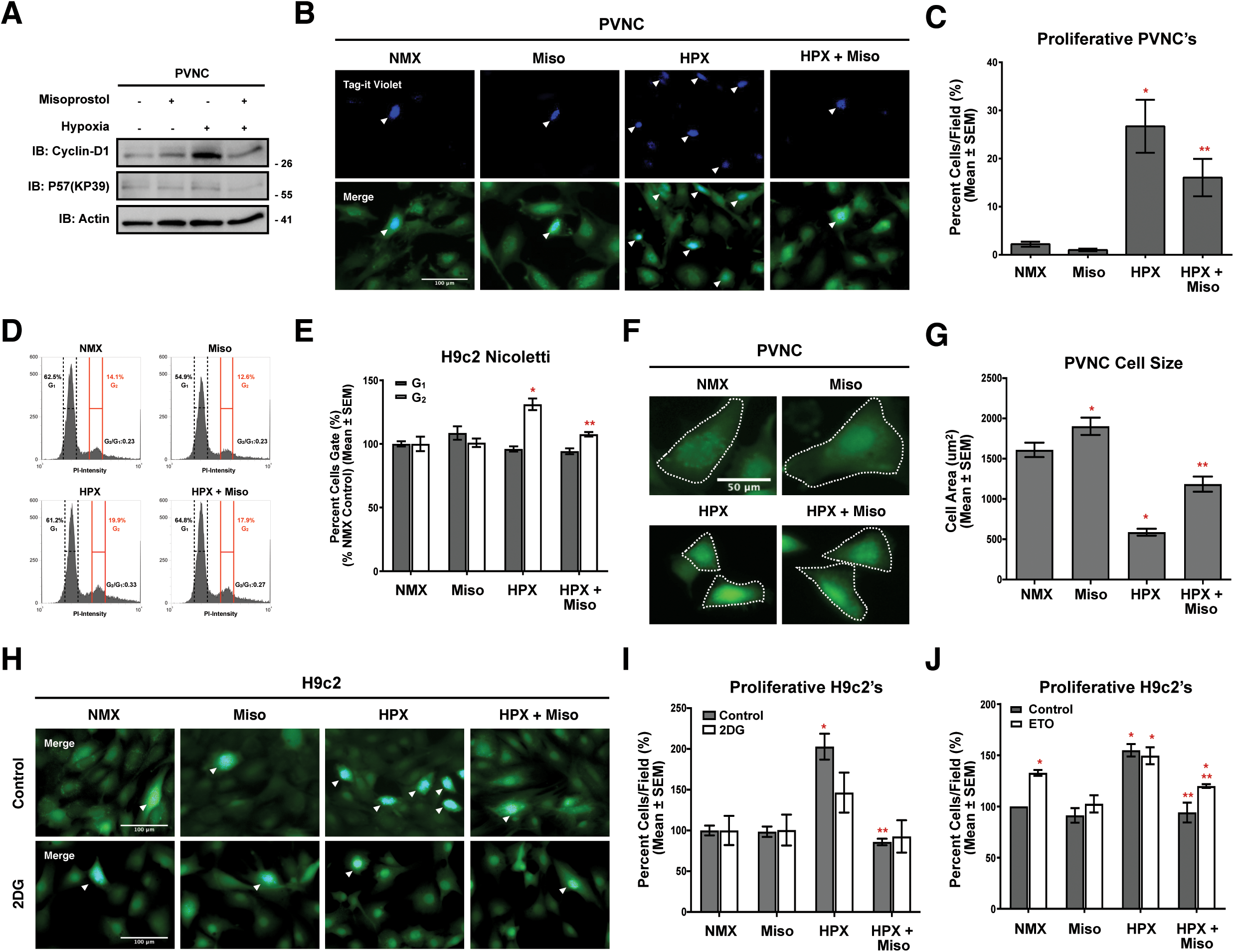
Misoprostol opposes hypoxia-induced neonatal cardiomyocyte proliferation. **(A)** Immunoblot for Cyclin-D1 and p57(KP39) in protein extracts from primary ventricular neonatal cardiomyocytes (PVNCs) treated with 10 μM misoprostol (Miso) ± 10% O_2_ (HPX) for 48 hours. **(B)** PVNCs treated as in (A) and stained with Tag-it Violet (blue) and calcein-AM (green) and imaged by standard fluorescence microscopy. **(C)** Quantification of Tag-it Violet positive cells in (B), where the number of blue cells is represented as a percentage (%) of the total number of (green) cells in 10 random fields. **(D)** Representative DNA Histograms of H9c2 cells treated as in (B), where G_1_ indicates cells that are 2N and G_2_ indicates cells that are 4N and G_2_/G_1_ ratio indicates the proliferative index. **(E)** Quantification of cells as in (D), comparing the percentage of cells gated into G_2_ vs. G_1_ across 6 independent experiments. **(F)** PVNCs treated as in (B) and stained with calcein-AM (green) and imaged by standard fluorescence microscopy. **(G)** Quantification of cells in (F), where cell surface area was calculated for >80 cells/condition in 10 random fields. **(H)** H9c2’s treated as in (A). 2 mM 2-Deoxy-D-glucose (2DG) was added to half of the conditions for 48 hours in order to inhibit glycolysis. Cells were stained with Tag-it Violet (blue) and calcein-AM (green) and imaged by standard fluorescence microscopy. **(I)** Quantification as in (C) of Tag-it Violet positive cells in (H) for 10 random fields. **(J)** Quantification of Tag-it Violet positive H9c2 cells treated as in (A), where 100 μM Etomoxir (ETO) was added to half of the conditions for 48 hours in order to inhibit fatty acid oxidation. The number of blue cells is represented as a percentage (%) of the total number of (green) cells in 10 random fields. All data are represented as mean ± S.E.M. *\*P*<0.05 compared with control, while *\*\*P*<0.05 compared with hypoxia treatment, determined by 1-way ANOVA or 2-way ANOVA where appropriate.

### Misoprostol opposes hypoxia-induced glycolytic flux

To further explore the role of glycolysis in proliferation, we monitored cytosolic NADH in H9c2 cells using the Peredox-mCherry biosensor [42], as an indicator of glycolytic activity. This ratiometric biosensor fluorescences red when in the presence of NAD^+^ and green in the presence of NADH. Exposure to hypoxia significantly increased the green fluorescent emission of Peredox-mCherry, indicating an increase in cytosolic NADH (Fig. 2A, -B). However, when cells were treated with misoprostol, the red emission was restored, suggesting that misoprostol inhibits hypoxia-induced NADH production. Next, we measured extracellular acidification (Seahorse XF-24) in H9c2 cells exposed to hypoxia and/or misoprostol treatment, and calculated glycolytic capacity and glycolytic reserve. Hypoxia exposure significantly increased extracellular acidification, and the calculated glycolytic capacity and glycolytic reserve (Fig.2C, -D, -E), which were attenuated by misoprostol treatment. Building on this, we moved to evaluate the accumulation of intracellular lactate, the end product of anaerobic glycolysis, using the FRET-based Laconic biosensor [43]. Shown in Figure 2F and -G, hypoxia exposure increased the FRET emission of the Laconic biosensor, indicating an increase in intracellular lactate accumulation, which was prevented in cells treated with misoprostol. This increase in lactate accumulation observed with hypoxia exposure corresponded with an increase in the expression of lactate dehydrogenase-A (LDHA; Fig.2H), which was also reduced with misoprostol treatment, suggesting that misoprostol-induced signaling alters myocyte gene expression to modulate glycolytic metabolism. We additionally evaluated intracellular pyruvate accumulation, a key metabolite in aerobic glycolysis, using the FRET-based Pyonic biosensor. Using this approach, we observed the reverse phenomenon to lactate accumulation, where hypoxia exposure decreased cellular pyruvate, which was restored to normal with misoprostol treatment (Fig.2I, -J). Finally, we determined that these results occurred in the absence of any observable changes in H9c2 glucose uptake (Fig.2K, -L). Taken together, these observations suggest that misoprostol can prevent hypoxia-induced anaerobic glycolytic flux through a mechanism that may involve alterations in gene expression, as is the case for LDHA.

**Figure 2.**
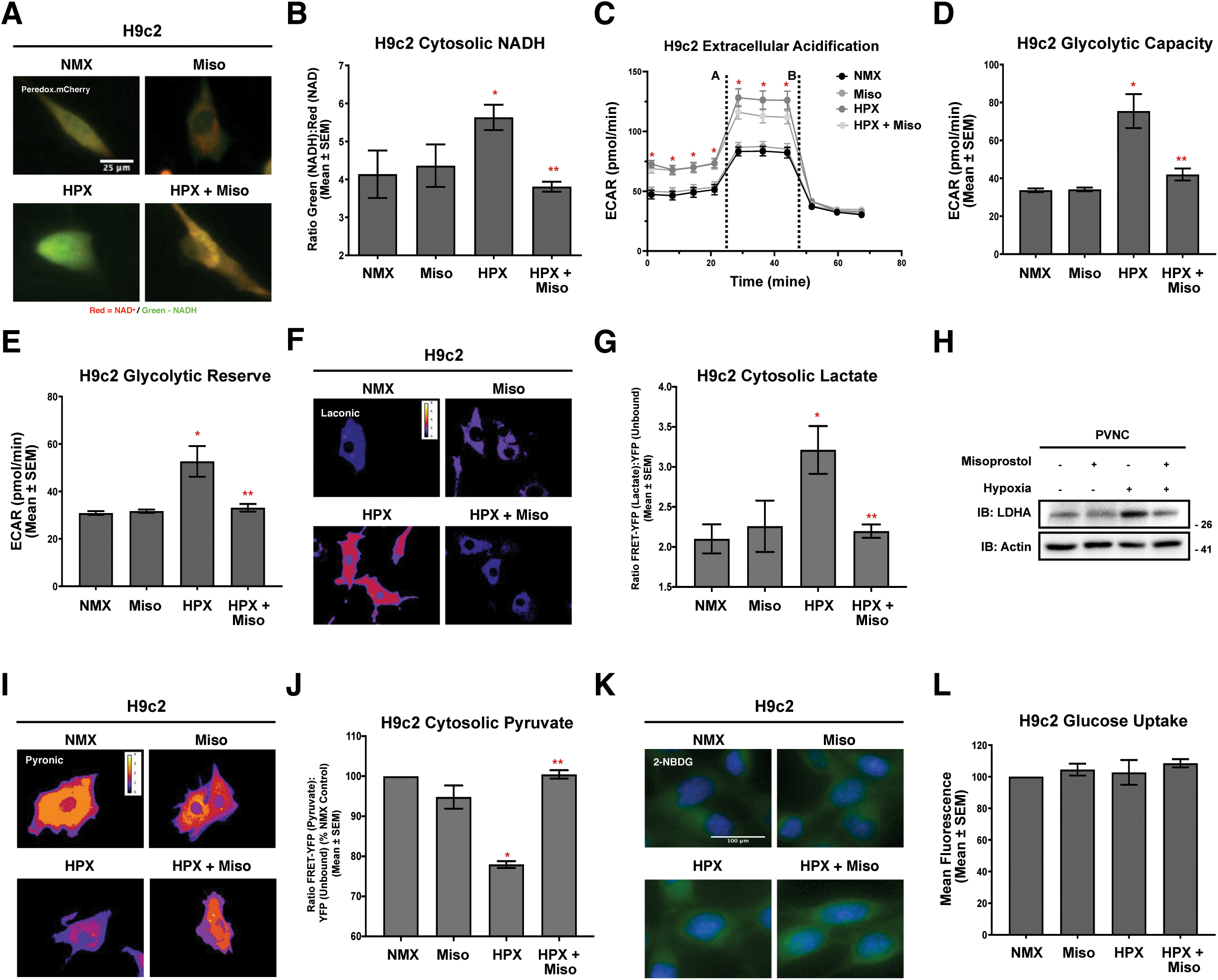
Misoprostol opposes hypoxia-induced glycolytic flux. **(A)** H9c2 cells treated with 10 μM misoprostol (Miso) ± 10% O_2_ (HPX) for 48 hours. Peredox-mCherry (where Red=NAD^+^, Green=NADH) was used to evaluate cytosolic reduced and oxidized NAD content in all conditions. Cells were imaged by standard fluorescence microscopy. **(B)** Quantification of cells in (A), where the ratio of green fluorescence (NADH) to red fluorescent signal (NAD^+^) in 15 random fields across 3 independent experiments. **(C)** H9c2 Extracellular Acidification Rate (ECAR) in cells treated as in (A) determined by Seahorse Extracellular Flux Analysis at baseline, following 1 μM oligomycin injection (a), and 50 mM 2DG injection (b) run across 3 independent experiments. **(D)** Quantification of H9c2 glycolytic capacity, calculated from (C). **(E)** Quantification of H9c2 glycolytic reserve, calculated from (C). **(F)** Ratiometric images of H9c2’s treated as in (A). Laconic was used to indicate cytosolic lactate content. Cells were imaged by FRET-based microscopy. **(G)** Quantification of cells in (F), where the FRET-YFP (lactate) signal was divided by the YFP (unbound biosensor) signal in 15 random fields across 3 independent experiments. **(I)** Ratiometric images of H9c2’s treated as in (A). Pyronic was used to indicate cytosolic pyruvate content. Cells were imaged by FRET-based microscopy. **(J)** Quantification of cells in (I), where the FRET-YFP (Pyruvate) signal was divided by the YFP (unbound biosensor) signal in 15 random fields across 3 independent experiments. **(K)** H9c2’s treated as in (A) stained with hoechst (blue) and 200 μM 2-NBDG (green) and imaged by standard fluorescence microscopy. **(L)** Quantification of cells in (K), where green fluorescent signal was normalized to cell area and quantified in 15 random fields across 3 independent experiments. All data are represented as mean ± S.E.M. *\*P*<0.05 compared with control, while *\*\*P*<0.05 compared with hypoxia treatment, determined by 1-way ANOVA or 2-way ANOVA where appropriate.

### Misoprostol promotes nuclear calcium accumulation and regulates gene expression

Given the central role of nuclear calcium signaling in the regulation of cardiac transcription factors, we first assessed nuclear calcium content using H9c2 cells and a nuclear targeted calcium biosensor (NLS-GECO) [41]. Although hypoxia and misoprostol alone had little impact on nuclear calcium content, when hypoxic cells were treated with misoprostol, we observed a significant increased nuclear calcium accumulation (Fig.3A,-B). Next, we evaluated how this calcium signal impacted the localization of the calcineurin-regulated transcription factor NFATc3, known to promote cardiomyocyte growth and maturation. Using H9c2 cells, we expressed a Myc-tagged NFATc3 construct (Myc-NFATc3), exposed cells to hypoxia, with and without misoprostol, and performed immunofluorescence using a primary Myc antibody and a fluorophore-conjugated secondary antibody. Consistent with our nuclear calcium data in Figure 3A, hypoxia and misoprostol on their own had little effect on the accumulation of NFATc3 in the nucleus, but the combination of hypoxia and misoprostol resulted in a more than 5-fold increase in nuclear NFATc3 accumulation (Fig.3C,-D). We confirmed this observation using cell fractionation studies in H9c2 cells that were exposed to 10% O_2_ with and without misoprostol for 48 hours. Here we observed that hypoxia and misoprostol decreased the cytosolic NFATc3 content and increased the nuclear content, without impacting the total, whole cell NFATc3 expression (Fig 3E). Next, we evaluated the role of calcineurin and NFATc3 on cytosolic NADH, as an indicator of hypoxia-induced glycolysis. Using the Peredox-mCherry biosensor, we treated cells with hypoxia and misoprostol, with and without the calcineurin inhibitor cyclosporine-A (CsA). Interestingly, CsA had little impact on hypoxia-induced NADH accumulation; however, the ability of misoprostol to block hypoxia-induced NADH accumulation was lost in the presence of CsA, suggesting that misoprostol works through calcineurin signaling to inhibit glycolysis (Fig3F, -G). As proof of concept experiments, we transfected cells with NFATc3, or an empty vector control, along with Peredox-mCherry, and exposed cells to hypoxia. We observed that hypoxia-induced NADH accumulation was prevented by NFATc3 expresion (Fig.3H), confirming that NFATc3 is sufficient to block hypoxia-induced glycolysis. In addition, we evaluated proliferation in cells expressing NFATc3, or an empty vector control, and observed less hypoxia-induced proliferation in cells expressing NFATc3 cells than in hypoxic control cells (Fig.I).

**Figure 3.**
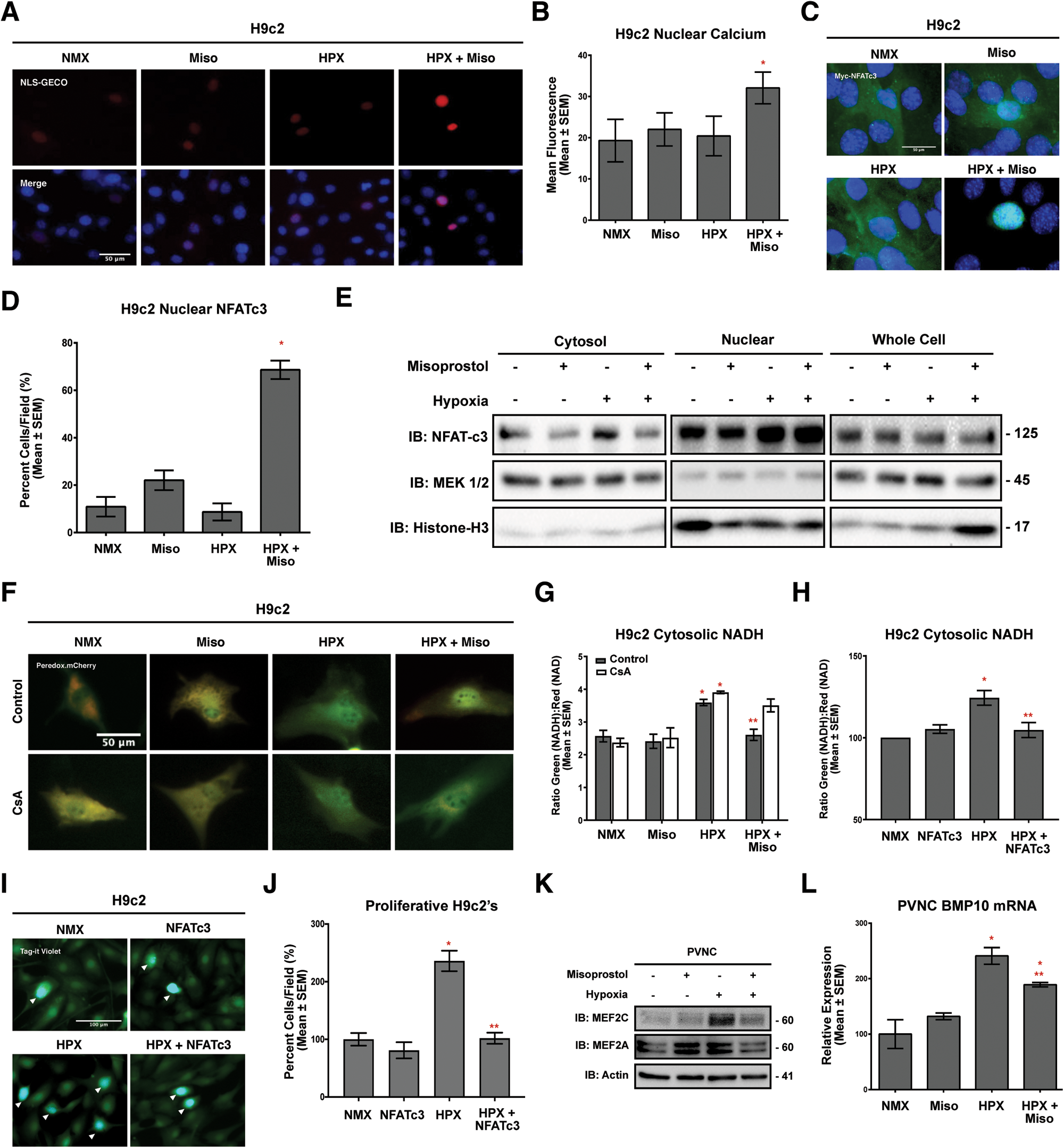
Misoprostol promotes nuclear calcium accumulation and regulates gene expression. **(A)** H9c2 cells treated with 10 μM misoprostol (Miso) ± 10% O_2_ (HPX) for 48 hours. NLS-R-GECO (red) was used to indicate nuclear calcium content in all conditions. Cells were stained with hoechst (blue) and imaged by standard fluorescence microscopy. **(B)** Quantification of cells in (A), where red fluorescent signal was normalized to cell area and quantified in 10 random fields. **(C)** H9c2 cells treated as in (A) and Myc-NFATc3 was transfected into each condition. Cells were fixed, stained with hoechst (blue), and immunofluorescence was performed using a Myc primary antibody (green). Cells were then imaged by standard fluorescence microscopy. **(D)** Quantification of cells in (C), where the number of cells with nuclear Myc-NFATc3, is presented as a percentage of the number of cells/field in 5 random fields. **(E)** H9c2 cells treated as in (A), Protein extracts were subjected to nuclear/cytosolic fractionation and were immunoblotted, as indicated. **(F)** H9c2’s treated as in (A). 2 μM CsA was added to half of the conditions for 48 hours in order to inhibit calcineurin-NFAT signalling. Peredox-mCherry (where Red=NAD^+^, Green=NADH) was used to indicate cytosolic reduced or oxidized NAD content in all conditions. Cells were imaged by standard fluorescence microscopy. **(G)** Quantification of cells in (F), where the ratio of green fluorescence (NADH) to red fluorescent signal (NAD^+^) in 15 random fields across 3 independent experiments. **(H)** Quantification of H9c2 cells treated with 10% O_2_ (HPX) for 48 hours and transfected pcDNA3 (control) or myc-NFATc3. Peredox-mCherry was used to indicate cytosolic NAD content. The ratio of green fluorescence (NADH) to red fluorescent signal (NAD^+^) was measured in >15 random fields across 6 independent experiments. **(I)** H9c2’s treated as in (H), cells were stained with Tag-it Violet (blue) and calcein-AM (green) and imaged by standard fluorescence microscopy. **(J)** Quantification of Tag-it Violet positive cells in (I), where the number of blue cells is represented as a percentage (%) of the total number of (green) cells in 10 random fields. **(K)** Immunoblot for MEF2C and MEF2A in protein extracts from PVNCs treated as in (A). **(L)** PVNCs treated as in (A). RNA was isolated and qRT-PCR was performed for BMP-10 expression. All data are represented as mean ± S.E.M. *P<0.05 compared with control, while *\*\*P*<0.05 compared with hypoxia treatment, determined by 1-way ANOVA or 2-way ANOVA where appropriate.

Next, we evaluated if misoprostol treatment could impact known cardiomyocyte proliferation pathways. Using western blot analysis of PVNCs we first observed that hypoxia increased the expression MEF2C, the MEF2 family member associated with fetal cardiomyocyte proliferation, while MEF2A expression remained unchanged from control (Fig.3J). Furthermore, the induction of MEF2C observed following hypoxia was prevented when combined with misoprostol treatment; however, MEF2A expression was sustained (Fig.3J). We also observed that hypoxia increased proliferative BMP10 expression by 58.5%, which was significantly reduced with the addition of misoprostol (Fig.3K).

### sNip opposes hypoxia-induced neonatal cardiomyocyte proliferation

Previously, we demonstrated that misoprostol treatment leads to the nuclear retention of the p65 subunit of NF-κB [20], and when combined with HIF1α activation, this led to the alternative splicing of Bnip3 in favour of sNip expression [20]. Thus, we performed western blot analysis on H9c2 cells exposed to hypoxia and/or misoprostol treatment to evaluate sNip expression. We observed that the combination of hypoxia and misoprostol increases the expression of the small Bnip3 isoform, sNip (Fig.4A). In order to define the role of sNip as a mechanism for misoprostol-induced blockade of cell cycle reentry, we performed a series of gain-of-function and loss-of-function experiments in PVNCs and H9c2 cells. When PVNCs were exposed to hypoxia, we observed a more than 3.8-fold increase in the number of proliferative nuclei, compared to normoxia-treated control cells. When these cells were transduced with a lentivirus delivering sNip, proliferation was reduced by 79.7%, back to control levels (Fig.4B, -C). In addition, we evaluated cell surface area in PVNCs exposed to hypoxia and transduced with sNip, and observed that sNip alone increased PVNCs surface area, and restored cell size in cells exposed to hypoxia (Fig.4D). Next, we performed a ‘rescue’ style experiment in H9c2 cells exposed to hypoxia and treated with misoprostol, where sNip function was reduced by a specific siRNA (Fig.4E, -F). Consistent with our observations in Figure 1, hypoxia induced cell proliferation, which was attenuated by misoprostol treatment. However, when cells were transfected with an siRNA targeting sNip, misoprostol treatment was no longer able to reduce cell proliferation (Fig. 4E, -F). In addition, we evaluated hypoxia-induced glycolysis in H9c2 cells expressing the Peredox-mCherry or the Laconic biosensors, along with sNip or empty vector control (Fig.4G, -H, -I). Consistent with the effect of misoprostol on glycolysis, sNip expression prevented the accumulation of both cytosolic NADH and intracellular lactate in cells exposed to hypoxia. Collectively, these observations support the notion that misoprostol exerts its effects on hypoxia-induced myocyte proliferation and glycolysis through the expression of sNip.

**Figure 4.**
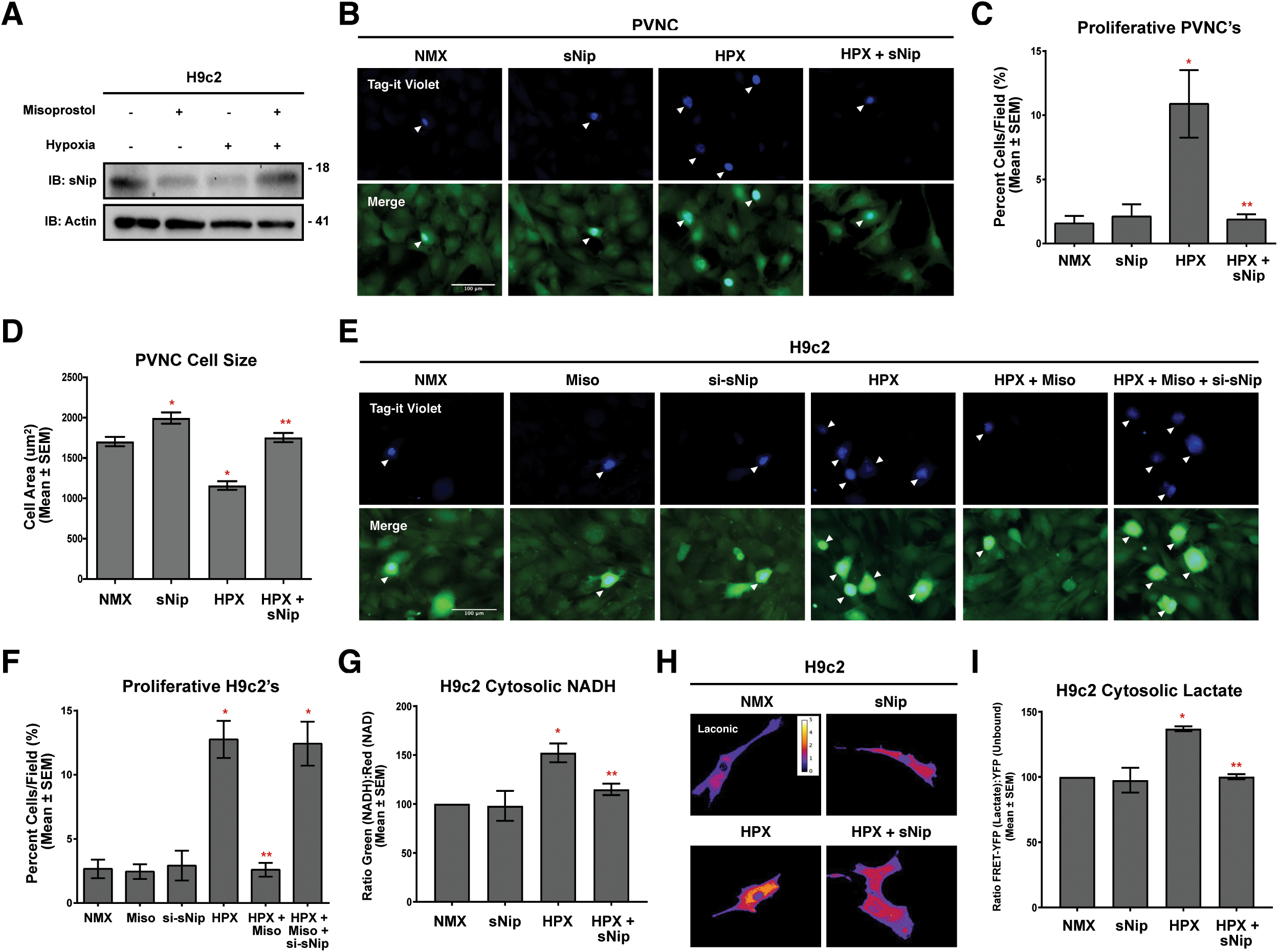
sNip opposes hypoxia-induced neonatal cardiomyocyte proliferation. **(A)** Immunoblot for sNip expression in H9c2 protein extracts treated with 10 μM misoprostol (Miso) ± 10% O_2_ (HPX) for 48 hours. **(B)** PVNCs treated with pLenti-HA-sNip ± 10% O_2_ (HPX) for 48 hours and stained with Tag-it Violet (blue) and calcein-AM (green) and imaged by standard fluorescence microscopy. **(C)** Quantification of Tag-it Violet positive cells in (B), where the number of blue cells is represented as a percentage (%) of the total number of (green) cells in 10 random fields. (D) Quantification of cells in (B), where cell surface area was calculated for >80 cells/condition in 10 random fields. **(E)** H9c2 cells treated as in (A), transfected with scrambled control si-RNA or si-sNip, and stained with Tag-it Violet (blue) and calcein-AM (green) and imaged by standard fluorescence microscopy. **(F)** Quantification of Tag-it Violet positive cells in (E), where the number of blue cells is represented as a percentage (%) of the total number of (green) cells in 10 random fields. **(G)** Quantification of H9c2 cells treated with 10% O_2_ (HPX) for 48 hours and transfected pcDNA3 (control) or HA-sNip. Peredox-mCherry was used to indicate cytosolic NAD content in all conditions. The ratio of green fluorescence (NADH) to red fluorescent signal (NAD^+^) was measured in >15 random fields across 3 independent experiments. **(H)** Ratiometric images of H9c2’s treated as in (G). Laconic was used to indicate cytosolic lactate content. Cells were imaged by FRET-based microscopy. **(I)** Quantification of cells in (H), where the FRET-YFP (lactate) signal was divided by the YFP (unbound biosensor) signal in 15 random fields across 3 independent experiments. All data are represented as mean ± S.E.M. *P<0.05 compared with control, while **P<0.05 compared with hypoxia treatment, determined by 1-way ANOVA or 2-way ANOVA where appropriate.

### sNip promotes nuclear calcium and NFATc3 accumulation

Next, we determined the impact of sNip expression on the calcium-dependent gene expression. First, we utilized the nuclear calcium biosensor NLS-GECO in H9c2 cells expressing sNip or empty vector, and exposed to normoxia or 10% O_2_ for 48 hours. We observed that both sNip on its own and in combination with hypoxia were sufficient to induce calcium accumulation in the nucleus (Fig.5A, -B).

**Figure 5.**
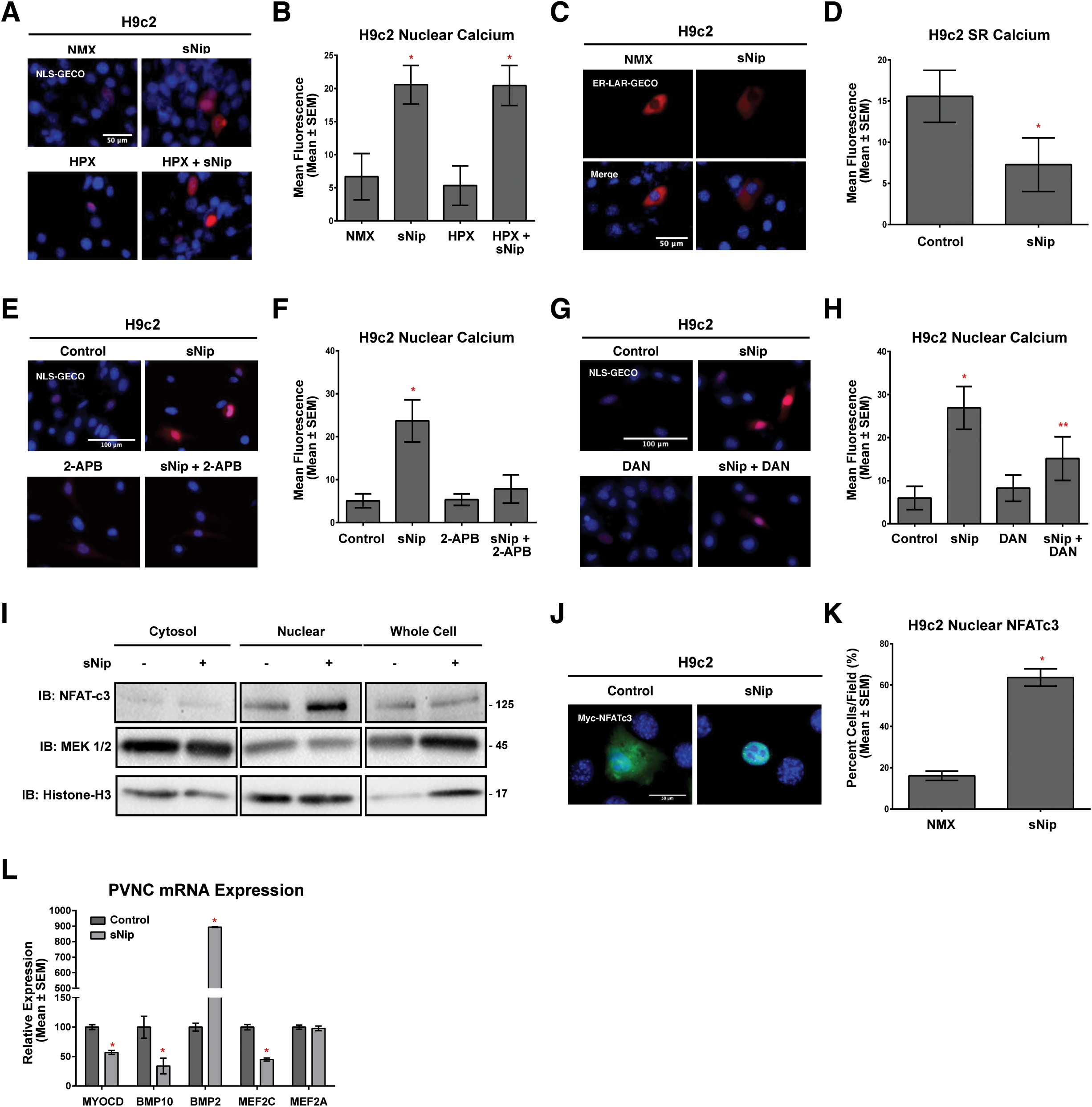
sNip promotes nuclear calcium and NFATc3 accumulation. **(A)** H9c2 cells transfected with sNip or empty vector control ± 10% O_2_ (HPX) for 48 hours. NLS-R-GECO (red) was used to indicate nuclear calcium content in all conditions. Cells were stained with hoechst (blue) and imaged by standard fluorescence microscopy. **(B)** Quantification of cells in (A), where red fluorescent signal was normalized to cell area and quantified in 10 random fields. **(C)** H9c2 cells transfected with sNip or empty vector control. ER-LAR-GECO (red) was used to indicate endoplasmic reticulum calcium content in all conditions. Cells were stained with hoechst (blue) and imaged by standard fluorescence microscopy. **(D)** Quantification of cells in (C) as in (B) across 10 random fields. **(E)** H9c2 cells transfected with sNip or empty vector control ± 2 μM 2-APB for 16 hours. NLS-R-GECO (red) was used to indicate nuclear calcium content in all conditions. Cells were stained with hoechst (blue) and imaged by standard fluorescence microscopy. **(F)** Quantification of cells in (E) as in (B) across 10 random fields. **(G)** H9c2 cells transfected with sNip or empty vector control ± 10 μM Dantrolene (DAN) for 16 hours, NLS-R-GECO (red) was used to indicate nuclear calcium content in all conditions. Cells were stained with hoechst (blue) and imaged by standard fluorescence microscopy. **(H)** Quantification of cells in (G) as in (B) across 10 random fields. **(I)** H9c2 cells treated as in (C), Protein extracts were subjected to nuclear/cytosolic fractionation and were immunoblotted, as indicated. **(J)** H9c2 cells treated as in (C). Myc-NFATc3 was included in all conditions to indicate transfected cells. Cells were fixed, stained with hoechst (blue), and probed for Myc (green) expression. Cells were then imaged by standard fluorescence microscopy. **(K)** Quantification of cells in (J), where the number of cells with nuclear Myc-NFATc3, is presented as a percentage of the number of cells/field in 5 random fields. **(L)** mRNA expression in PVNCs transduced with scrambled control virus or pLenti-sNip. RNA was isolated and qRT-PCR was performed for Myocardin (MYOCD), BMP-10, BMP-2, MEF2C and MEF2A expression. All data are represented as mean ± S.E.M. *\*P*<0.05 compared with control, while *\*\*P*<0.05 compared with hypoxia treatment, determined by 1-way ANOVA or multiple T-Tests where appropriate.

Given that the endoplasmic/sarcoplasmic reticulum (ER/SR) contains a large pool of intracellular calcium and is continuous with the outer membrane of the perinuclear envelope, we sought to determine if the ER/SR was the source of cellular calcium that was accumulating in the nucleus under the influence of sNip. We used the ER targeted calcium biosensor (ER-LAR-GECO), described previously [20,22,41], to assess ER/SR calcium content. Interestingly, when expressed in H9c2 cells sNip reduced steady-state ER calcium content by 58.8% compared to control (Fig. 5C, -D). In a parallel series of experiments, sNip-induced nuclear calcium accumulation was inhibited when cells were treated with the IP_3_ receptor inhibitor, 2-APB [47](Fig.5E, -F), or the ryanodine receptor antagonist dantrolene, where the magnitude of this blockade was greater for 2APB (Fig.5G, -H).

Next, we expressed sNip in H9c2 cells to evaluate its effect on transcription factor localization and gene expression. Using subcellular fractionation to separate components of the cytosolic compartment from components of the nuclear compartment, we observed that sNip increased the nuclear localization of NFATc3, without impacting total cell NFATc3 (Fig.5I). We confirmed this observation using immunofluorescence, where sNip expression resulted in a 2.96-fold increase in the nuclear localization of NFATc3 (Fig.5J, -K). In addition, sNip transduction in PVNCs reduced the expression of myocardin (43%), BMP10 (66%) and MEF2C (55%), and increased the expression of the NFAT target gene BMP2 by 794% (Fig.5L), determined by real-time PCR.

### Misoprostol prevents hypoxia-induced cardiomyocyte proliferation in vivo

To evaluate our observation *in vivo*, we used a rodent model of neonatal hypoxia, where neonatal Long-Evans rats are exposed to 10% oxygen, and treated with misoprostol or vehicle control for 5-days. Importantly, we did not observe any acute adverse effects associated with misoprostol treatment in hypoxic animals. Histological examination of the neonatal left ventricle revealed an increase in the number of myo-nuclei by an average of 9.5 nuclei per field, which was prevented by misoprostol treatment (Fig.6A, -B). Interestingly, this degree of hypoxia was not sufficient to induce apoptosis determined by TUNEL staining, or cardiac fibrosis determined by Trichrome staining (Fig.6A). However, immunofluorescence staining using a primary antibody targeting the specific proliferation marker PHH3, identified an increased number of proliferative nuclei in hypoxia exposed hearts, which was completely eliminated in misoprostol treated hearts (Fig.6A, -C). In addition, we confirmed that the combination of hypoxia and misoprostol treated increased sNip expression *in vivo* (Fig.6D), consistent with our observations in cultured myoblasts.

**Figure 6.**
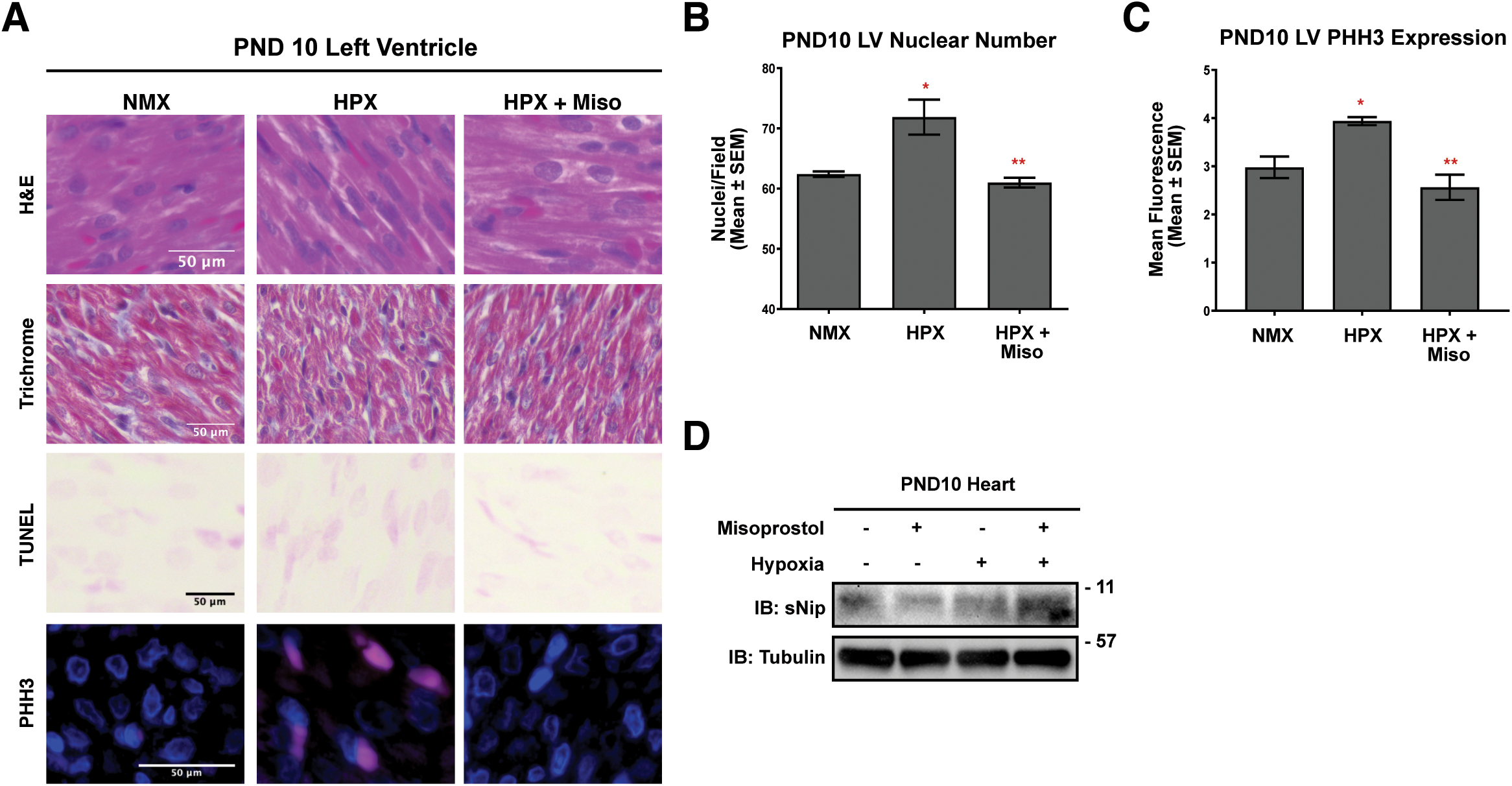
Misoprostol prevents hypoxia-induced cardiomyocyte proliferation *in vivo*. **(A)** representative images of Hematoxylin and Eosin (H&E), Masson’s Trichrome, and TUNEL staining, as well as immunofluorescence for Phospho-Histone H3 (PHH3) in the left ventricle of PND10 rat pups exposed to hypoxia (10% O_2_) ± 10 μg/kg misoprostol for 5 days. **(B)** Quantification of left ventricular nuclei number in the PND10 rat heart in (A) (n=3 animals/condition, with 10 fields/animal). (C) Quantification of left ventricular PHH3 expression PND10 rat heart in (A) (n=2 animals/condition, with 10 fields/animal). **(D)** Immunoblot for sNip expression in protein extracts from whole PND10 hearts treated as in (A). All data are represented as mean ± S.E.M. *\*P*<0.05 compared with control, while *\*\*P*<0.05 compared with hypoxia treatment, determined by 1-way ANOVA or 2-way ANOVA where appropriate.

## Discussion

Neonatal hypoxic injury is considered detrimental to cardiac development, increasing the risk of pediatric heart failure, and lifelong cardiovascular abnormalities. However, recently it has been observed that the underlying cause may involve aberrant cardiomyocyte proliferation. In this report, we show that hypoxia drives cardiomyocyte proliferation, concurrent with activation of a MEF2C and BMP10 pathway, and enhanced glycolytic metabolism. We also provide evidence that PGE1 signalling through misoprostol treatment is able to activate a secondary and opposing pathway, enhancing Bnip3 splicing to produce sNip. This alternative splicing of Bnip3 results in nuclear calcium accumulation, NFATc3 activation, inhibition of glycolysis, and attenuation hypoxia-induced proliferation. These findings are consistent with our previous results demonstrating that misoprostol treatment restores mitochondrial respiration in hypoxic cardiomyocytes [20].

The results in the present study expand on previous reports that early life hypoxia exposure is sufficient to drive neonatal cardiomyocyte proliferation. These previous studies have mainly focused on the regenerative capacity of hypoxia-induced cardiomyocyte proliferation (20, 32, 34). Interestingly, maintaining a capacity for proliferation after birth appears to depend on the metabolic status of the cardiomyocyte, where the switch from glycolysis to oxidative phosphorylation and the resultant accumulation of cytotoxic free radicals is thought to result in cardiomyocyte senescence [11]. Moreover, recent evidence further suggests that preventing this metabolic shift is key to maintaining c-kit^+^ cardiac progenitor cells in culture [48].

While the literature to date has primarily focused on the role of cellular metabolism, our data provides insight into the underlying hypoxia-driven mechanisms promoting neonatal cardiomyocyte proliferation. In this report, we confirm that expression of MEF2C is hypoxia inducible, and that this may be a key cardiac-enriched initiator of a hypoxia-induced cardiomyocyte proliferation [18,19]. Furthermore, evidence supports the existence of a feed-forward loop promoting proliferation that hinges on MEF2C-dependent activation of myocardin and downstream BMP10 expression. Importantly, loss of the myocardin-BMP10 signalling pathway negatively impacts cardiomyocyte proliferation [21]. BMP10 also appears to be required for maintaining MEF2C expression and preventing myocyte maturation, which may function to propagate proliferation and alter the structure of the developing heart [49].

A number of previous studies have assessed methods to extend the period of post-natal cardiomyocyte proliferation [7–9,11]. However, in the contexts of early life hypoxic/cyanotic events and cardiac regeneration, there may be direct benefits to pharmacologically targeting cardiomyocyte proliferation. Our data demonstrates that elevated sNip expression through misoprostol treatment can inhibit the MEF2C-Myocardin-BMP10 pathway, and simultaneously increase BMP2 expression. Previous work from our group, and others, have demonstrated that while sNip is hypoxia-inducible, NF-κB activity is crucial for its expression in cardiomyocytes [20,38]. Interestingly, much of sNip’s opposition to cardiomyocyte proliferation appears to be based on its ability to promote the transfer of calcium from the SR/ER to the nucleus. This builds on our recent findings that sNip possesses a C-terminal ER retention signal, and that unlike full-length Bnip3, sNip expression in cardiomyocytes functions to sustain nuclear calcium accumulation [20]. Nuclear calcium signaling is known to activate calcium-calmodulin dependent transcriptional regulators, such as the calcineurin-NFAT pathway [23–25,27]. Our findings demonstrate that NFATc3 is sufficient to oppose both hypoxia-induced glycolysis and proliferation in cultured myocytes. These observations are consistent with previous work demonstrating that NFAT and NF-κB are considerably less efficient at producing a hypertrophic response alone than when the two are expressed together [50].

Collectively, our data demonstrates that the activation of PGE1 signalling through misoprostol treatment is sufficient to mitigate hypoxia-induced proliferation in neonatal cardiomyocytes through a mechanism involving alternative splicing of Bnip3, activation of NFATc3, and inhibition of hypoxia-induced glycolysis. Moreover, our findings suggest that pharmacological fine-tuning of the underlying molecular mechanisms that control hypoxia-induced cardiomyocyte proliferation may represent a novel strategy to prevent pediatric cardiac dysfunction, reduce the risk of early-life heart failure, and have implications for cardiac regenerative medicine.

## Acknowledgments

This work was supported by a Natural Sciences and Engineering Research Council (NSERC) Canada Discovery Grant and a Heart and Stroke Foundation of Canada Grant-in-Aid to J.W.G. T.L.I. is funded through the SFRG and UCRP programs from the University of Manitoba. J.W.G. and A.R.W. are members of the DEVOTION Research Cluster. M.D.M., J.T.F., N.S. and C.D. are supported by studentships from the Children’s Hospital Foundation of Manitoba and Research Manitoba, M.D.M. received support from the DEVOTION research cluster.

## Author Contributions

M.D.M. and J.W.G. conceived and coordinated the study. M.D.M. and J.W.G. wrote the paper. M.D.M. designed and conducted most of the experiments. M.D.M. was also responsible for data analysis and presentation. J.T.F. conducted flow cytometry and fractionation experiments. D.C. designed and conducted the qRT-PCR experiments. N.S. designed and conducted Seahorse XF-24 experiments and data analysis. C.D. conducted immunofluorescence experiments in PND10 heart tissue. T.L.I. designed and conducted the in vivo hypoxia experiments. J.W.G., T.L.I. and A.R.W. were responsible for funding acquisition. All authors reviewed the results, edited, and approved the final version of the manuscript.

